# Presenilin-mediated cleavage of APP regulates synaptotagmin-7 and presynaptic plasticity

**DOI:** 10.1101/258335

**Authors:** Gaël Barthet, Tomàs Jordà-Siquier, Julie Rumi-Masante, Fanny Bernadou, Ulrike Müller, Christophe Mulle

## Abstract

Presenilin (PS), the catalytic subunit of γ-secretase and its main substrate the amyloid precursor protein (APP) are mutated in a large majority of patients with familial Alzheimer disease. PS and APP interact with proteins of the neurotransmitter release machinery but the functional consequences of these interactions are unknown. Here we report that genetic deletion of presynaptic PS markedly decreases the axonal expression of the Ca^2+^ sensor synaptotagmin-7 (Syt7), and impairs synaptic facilitation and replenishment of release-competent synaptic vesicles. These properties are fully restored by presynaptic re-expression of Syt7. The regulation of Syt7 expression occurs post-transcriptionally and depends on γ-secretase activity. In the combined absence of both APP and PS1, the loss of Syt7 is prevented, indicating that the action of γ-secretase on presynaptic mechanisms depends on its substrate APP. The molecular mechanism involves the substrate of PS, APP-βCterminal (APP-βCTF), which interacts with Syt7 and accumulates in synaptic terminals under conditions of pharmacological or genetic inhibition of γ-secretase. These results reveal a role of PS in presynaptic mechanisms through regulation of Syt7 by APP-dependent cleavage, and highlight aberrant synaptic vesicle processing as a possible new pathway in AD.

## INTRODUCTION

The etiology of Alzheimer disease (AD) is complex. The familial forms of Alzheimer disease (FAD) are caused by mutations of two presenilin (PS) paralogs PS1 or PS2^1^ or of the amyloid precursor protein (APP)^2^ indicating a central role of these genes in the disease. PS is the catalytic subunit of the intramembrane protease γ-secretase which cleaves several type 1 transmembrane proteins including APP^3^. The involvement of both PS and APP in the same enzymatic reaction indicates a possible role for APP fragments processed by PS in AD etiology. Within the amyloidogenic pathway, APP is first cleaved by β-secretase (BACE1) producing the APP βCterminal fragment (APP-βCTF), a transmembrane stub further processed through the intramembrane proteolytic activity of γ-secretase which releases Aβ peptides and the APP-intracellular domain (AICD). While molecular studies have mostly focused on Aβ, possible roles for all three APP fragments in AD have been reported^4^. At the clinical level, synaptic dysfunction and loss are the leading pathological correlates of cognitive impairment in AD^5–7^ suggesting a role for AD-related genes in synaptic transmission.

Accordingly, the roles of both PS and APP in synaptic transmission and plasticity have been studied in several cellular and mouse models (reviewed in ref. ^8^ for PS and in ref. ^9^ for APP). Both proteins are abundantly expressed in the presynaptic active zone^10–12^ where they interact with proteins of the synaptic release machinery ^13–15^. Therefore, the physiological role of both PS and APP in presynaptic mechanisms through interactions with proteins of the presynaptic release machinery needs to be investigated. To address this, we investigated the presynaptic function of PS and APP at hippocampal mossy fiber (Mf) synapses formed between the dentate gyrus (DG) and CA3 pyramidal cells (Mf-CA3 synapses). Mf-CA3 synapses are large synapses with multiple release sites and are characterized by a large dynamic range of presynaptic plasticity^16^. Moreover, Mfs are endowed with a prominent expression of γ-secretase both in rodents^17^ and in humans^18^. We identified Syt7 as a key presynaptic actor linking the absence of PS or inhibition of γ-secretase activity to presynaptic accumulation of APP-βCTF and impaired presynaptic mechanisms.

## Results

### An optogenetic strategy to combine cell-specific activation with gene deletion in DG granule cells

We have developed optogenetic tools that combine cell specific activation with cell specific gene deletion of DG granule cells (GCs) to study the role of PS at Mf-CA3 synapses. The expression of the C1ql2 gene is largely restricted to the DG in the hippocampus^19^. We cloned a sequence of the promoter rich in CpG islands which includes a 1 kb fragment upstream of the transcriptional start site and part of the 5’ untranslated region of the mRNA sequence to create a C1ql2 minimal promoter (**Supplementary Fig. 1a**). The minimal promoter was used in a lentiviral bicistronic construct to control the expression of both ChIEF (a variant of ChR2, see ref. ^20^) fused to the red fluorescent protein tomato and the Cre recombinase, both cDNAs being separated by a 2A sequence (**Fig. 1a**). A similar strategy combining expression of ChIEF and Cre was recently applied using an internal ribosomal entry site (IRES) sequence^21^. In brain sections from mice injected with this construct, ChIEFtom is selectively expressed in the DG (**Fig. 1b**). Moreover, an extensive study of C1ql2-mediated expression of reporter genes validated the bicistronic 2A strategy, the efficiency of the method and the specific targeting of mature Mfs (**Supplementary Fig. 1b-e**). We used this tool in PS1floxed/PS2KO mice to characterize the properties of Mf-CA3 synapses lacking PS presynaptically, but still expressing PS1 postsynaptically (**Fig. 1c**).

**Figure 1.**
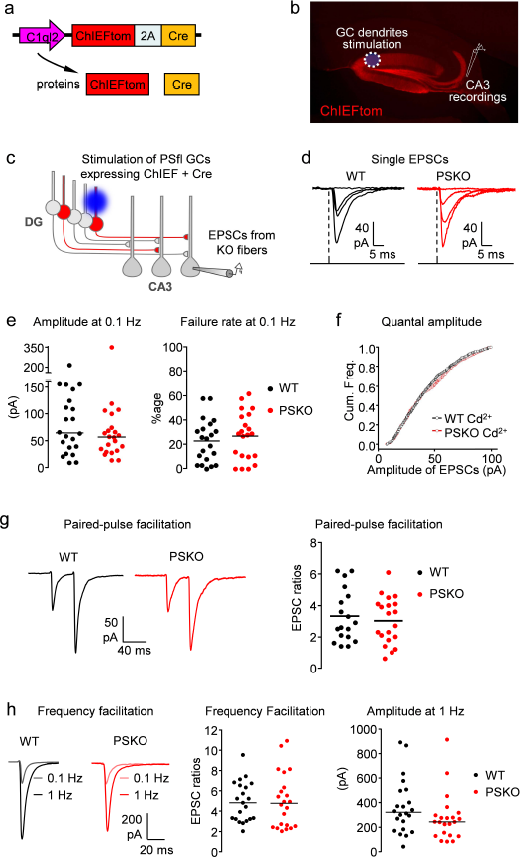
Study of PS presynaptic function at Mf/CA3 synapses using an optogenetic strategy. (**a**) Scheme of lentiviral construct which co-expresses ChIEF fused to the fluorescent protein tomato and the Cre recombinase. The promoter C1ql2 restricts expression to DG granule cells (GCs). (**b**) Picture of a parasagittal section of a mouse brain injected with a virus coding for ChIEFtom. (**c**) Strategy to selectively study Mf/CA3 synapses that are PSKO presynaptically and intact postsynaptically. The blue light triggers the firing of granule cells that express ChIEF and by obligation (bicistronic construct) the Cre recombinase. In PS *“floxed”* conditional knockout mice, the activated GCs do not express PS; EPSCs evoked in CA3 neurons by light pulses in the DG arise from presynaptic PSKO Mf terminals. (**d**) Representative traces of CA3 EPSCs evoked by single light-pulses focused as in “B”. (**e**) Basal transmission at Mf/CA3 synapses is not altered by PS genotype. *Left*, scatter plots with medians of EPSCs amplitude recorded in CA3 neurons at 0.1 Hz in WT or PSKO condition (n=22/genotype). *Right*, scatter plots with mean of failure rate. (**f**) Cumulative frequencies of EPSCs amplitude in presence of Cd^2+^. (**g**) *Left*, representative traces of paired-pulses EPSCs evoked by two light-pulses separated by 40 ms. *Right*, analysis of several recordings obtained as in “Left”. The facilitation is represented by the amplitude of the second EPSC normalized to the amplitude of the first EPSC. (**h**) *Left*, representative traces of EPSCs evoked by light-pulses at 0.1 or 1 Hz. *Right*, analysis of several recordings obtained as in “Left”. The frequency facilitation is represented by the amplitude of EPSCs obtained at 1 Hz normalized to the amplitude of EPSCs obtained at 0.1 Hz.

We first recorded from ChIEF-expressing GCs in the current-clamp mode, and observed that a 1 ms light-pulse was sufficient to trigger an action potential (AP) (**Supplementary Fig. 1f**). Trains of light pulses at 20 Hz reliably triggered trains of APs up to several seconds without failures (**Supplementary Fig. 1f**). By illuminating the dendritic layer of the DG, we could activate a restricted population of GCs, and consequently a restricted number of Mfs (**Fig. 1b,c** and **Supplementary Fig. 1g**). Mf-EPSCs evoked in CA3 pyramidal cells using this stimulation protocol displayed properties comparable to those of Mf-EPSCs evoked by minimal electrical stimulation^22^. Mean EPSC amplitude, failure rate and short-term plasticity, including paired-pulse facilitation and low frequency facilitation (at 1 Hz) were comparable (**Supplementary Fig. 1h**) between the two modes of stimulation.

### Loss of presynaptic PS impairs facilitation and SV replenishment

We virally transduced the C1ql2-ChIEFtom-2A-Cre construct into PS1floxed/PS2-KO mice (PS1 conditional KO mice backcrossed to constitutive PS2^-/-^ mice). This allowed the recording of Mf-EPSCs from presynaptic PSKO Mf-CA3 synapses when the dendrites of Cre-expressing GCs are stimulated by light (**Fig. 1c**). We compared the properties of EPSCs at WT (ChIEFtom2A-Cre expression in WT mice) and PSKO Mf-CA3 synapses (**Fig. 1d**). The mean basal amplitude of Mf-EPSCs and the failure rate were comparable between the two genotypes when GCs were stimulated at low frequency (0.1 Hz), suggesting similar initial release probability (**Fig. 1d,e**). To further study basal synaptic transmission, we recorded Mf-EPSCs in the presence of 32 μM of Cd^2+^ (a bivalent cation that blocks Ca^2+^ channels), leading to partial inhibition of transmitter release at this concentration (**Supplementary Fig. 1i**) Under these conditions, 95% of Mf-EPSCs at 1 Hz stimulation were estimated to be due to the quantal release of glutamate (see Methods). No difference in the quantal amplitude of Mf-EPSCs between the two genotypes was observed (**Fig. 1f,** mean: WT=42.4 pA; PSKO=43.8 pA / median: WT=PSKO=38.0 pA).

We then compared short-term plasticity between the two genotypes. Paired-pulse facilitation (40 ms interval) and frequency facilitation, observed when switching the frequency of stimulation from 0.1 to 1 Hz, did not differ between WT and PSKO Mf-CA3 synapses (**Fig. 1g,h**). In contrast, the large facilitation triggered by short trains of light stimulation of GC dendrites at 20 Hz (train facilitation) was markedly smaller at PSKO Mf-CA3 synapses (**Fig. 2a**). Interestingly, the deletion of presynaptic PS1 alone was sufficient to cause a deficit in train facilitation (**Supplementary Fig. 2a**). The deficit in train facilitation cannot be attributed to changes in initial release probability as indicated above. PS inactivation may directly impact on the excitability of DG GCs. However we did not observe any difference in spiking activity of GCs in response to trains of light stimulation in GC dendrites. The amplitude or half-width of action potentials recorded in DG GCs along the 20 Hz train did not differ between WT and PSKO GCs (**Supplementary Fig. 2b**). Overall, inactivation of PS at Mf-CA3 synapses neither affected release probability nor quantal amplitude under basal conditions, but strongly decreased facilitation induced by trains of Mf stimulation by affecting short-term plasticity mechanisms.

**Figure 2.**
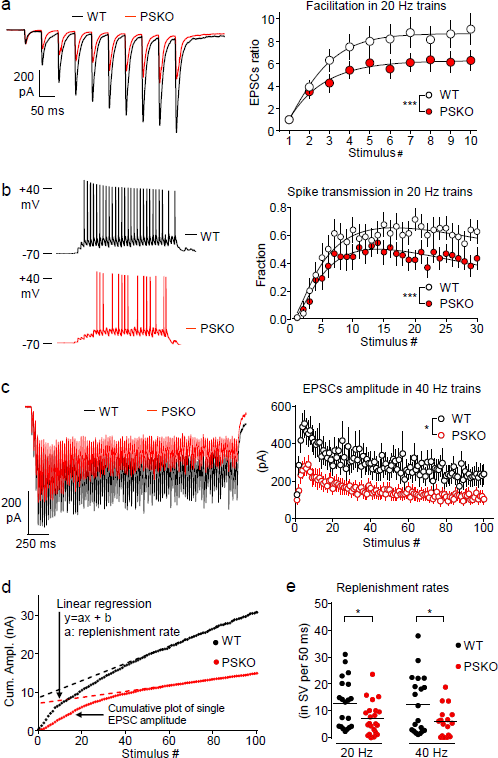
PS promotes presynaptic facilitation, spike transmission and SV replenishment. (**a**) *Left*, representative traces of EPSCs evoked by 10 repetitive light-pulses at 20 Hz. *Right*, analysis of several recordings obtained as in *“Left”.* The facilitation is represented by the amplitude of EPSCs normalized to the amplitude of the first EPSC. The facilitation is strongly impaired by absence of presynaptic PS. Two-way ANOVA, n=19 in WT, 16 in PSKO, *p=0.0004.* (**b**) *Left*, representative traces of CA3 spikes in response to 30 stimulations at 20 Hz. *Right*, plot of the fractions of spike transmission (for 16 recorded neurons per genotype; 5 sweeps per neurons) in WT or PSKO genotype. The highest rate of transmission (over 50 percent) is reached around 15 stimuli and the plot in PSKO is lower than in WT condition (Two-way ANOVA, *p<0.0001).* (**c**) *Left*, representative EPSCs evoked by 100 stimuli at 40 Hz. After the initial rising phase (facilitation), the amplitude drops and reaches a steady state that is reduced by absence of PS. (**d**) Graph of the cumulative amplitude of Mf-EPSCs in WT or PSKO condition from trains of stimulation plotted against the stimulus number. A linear regression fits the second part of the cumulative plot where a steady state is reached. The slope is proportional of the replenishment rate. (**e**) Scatter plots with means of replenishment rates calculated neuron per neuron are represented by genotype and frequency. Absence of PS decreases the rates of replenishment (unpaired t-test p<0.05; n=20 in WT, 22 in PSKO).

The broad dynamic range of EPSC amplitudes from failures (0.1 Hz) to large currents in response to high-frequency bursts of stimulation represents a high-pass filter for the transfer of information at Mf-CA3 synapses. We explored the impact of the short-term plasticity deficit on spike transfer at DG-CA3 connections by performing current-clamp recordings of CA3 pyramidal cells while stimulating GC dendrites with light. In minimal stimulation conditions, the amplitude of Mf-EPSPs at low stimulation frequencies (0.1 to 1 Hz) is small and only occasionally triggered spike discharge in CA3 PCs (**Supplementary Fig. 2c**). Spike transmission increased dramatically in response to trains of stimulation at 20-40 Hz (**Fig. 2b** and **Supplementary 2d**) confirming the role of Mf-CA3 synapses as high-pass filters for information transfer within the hippocampal circuit^23^. Within a train of 30 stimuli, the fraction of spike transmission at WT Mf-CA3 synapses increased from 0 to 0.63 after 15 stimuli and then slowly decreased (**Fig. 2d**). In the absence of PS, spike transfer at DG-CA3 connections was markedly less efficient reaching on average a fraction of 0.45 spikes per stimulus after 15 stimuli (**Fig. 2b**). Altogether, these results indicate that PS is essential for presynaptic facilitation that supports high-pass filtering of spike transfer at DG-CA3 connections

Synaptic transmission depends on the priming of release-competent synaptic vesicles (SVs), a mechanism dynamically regulated by the replenishment of the readily-releasable pool of SVs^24^. Because inactivation of PS impacts Mf-CA3 synapses under conditions of trains of presynaptic stimulation, we postulated that the replenishment of release-competent SVs could be impaired. To test this hypothesis, we used long trains of stimulation at 20-40 Hz that involve the recycling pool of SVs (**Fig. 2c** and **Supplementary Fig. 2e**). We estimated the replenishment rate of SVs at Mf-CA3 synapses by plotting the cumulative amplitude of Mf-EPSCs against stimulation number^25^ (**Fig. 2d**). The rate of replenishment was estimated for each recorded CA3 cell by fitting the linear part of the curve for 100 stimuli trains at 20 Hz by linear regression (**Fig. 2d),** and taking into account the average quantal size measured above (**Fig. 1f**). The replenishment rate at Mf-CA3 synapses was reduced approximately by half in PSKO conditions as compared to WT conditions (WT and PSKO values, from 13 to 7 SVs per 50 ms, **Fig. 2e**).

### Unaltered presynaptic terminals

We investigated whether the reduced facilitation and impaired replenishment rate of SVs resulted from the degeneration of presynaptic Mf terminals lacking PS, possibly leading to smaller Mf boutons with fewer release sites. To investigate this possibility, we studied the morphology of Mf boutons expressing membrane-targeted YFP as a morphological fluorescent marker^26^, by confocal imaging followed by 3D reconstruction. In both genotypes, Mf boutons had an average volume of 16 μm^3^ (on 200 Mf boutons, **Supplementary Fig. 2f**). We then stained Mf boutons for Bassoon, a presynaptic marker for release sites and we performed stimulated-emission-depletion (STED) microscopy to evaluate differences in the number of Bassoon puncta between WT and PSKO Mf boutons. We did not observe any difference in the number (16.7 in WT; 17.0 in PSKO) and in the area (0.073 μm^2^ in WT; 0.078 μm^2^ in PSKO, n=82 Mf boutons from 3 mice per genotype) of Bassoon-positive puncta (**Supplementary Fig. 2g**) suggesting that impaired facilitation at Mf-CA3 synapses in PSKO does not result from fewer active zones. These morphological data indicate that the impairment of presynaptic function in PSKO is not caused by a broad alteration of the structure or composition of Mf terminals but more likely results from a distinct alteration at the molecular level.

### Marked drop of Syt7 axonal expression in the absence of PS

Recent evidence indicates that the Ca^2+^ sensor synaptotagmin 7 (Syt7) is necessary for both presynaptic facilitation^27^ and replenishment of release-competent SV^28^, two presynaptic mechanisms being impaired in the absence of PS. We tested whether the functional impairment of PSKO Mf-CA3 synapses results from a decreased expression of Syt7 by immunolabelling Syt7 in mouse brain slices. We observed that Syt7 is an abundant protein in the mouse brain (**Supplementary Fig. 3a-c),** and is enriched in the *stratum lucidum* (SL) where Mfs make synaptic contacts with CA3 pyramidal cells, in agreement with its role in synaptic facilitation^27^. Notably, expression of Syt7 in the *SL* contrasts with the adjacent *stratum radiatum (SR)* where recurrent CA3/CA3 inputs make synapses devoid of marked presynaptic facilitation (**Fig. 3a**). In brain slices in which PS is absent only in DG-GCs, the labelling of Syt7 in the *SL* was dramatically decreased, indicating a loss of Syt7 in PSKO Mf axons (**Fig. 3a**). We measured the mean pixel intensity of staining in the *SL* and normalized it to the mean pixel intensity in the SR, a region not genetically manipulated (**Fig. 3b**). We observed an average decrease of 40% (**Fig. 3c),** a result likely minimized by the infection rate of GCs by the Cre-expressing lentivirus (transduction in around 50% of GCs). We then prepared protein lysates from laser capture microdissections of the *SL* and hilus to quantify the expression level of the different Syt7 splice variants. The Syt7 antibody labeled two bands at 46 and 80 kDa (**Fig. 3d),** corresponding to different splice variants^29^. The shorter variant carries an Nterminal transmembrane region followed by two Cterminal Ca^2+^ binding domains (C2A and C2B)^30^ and has been shown to rescue presynaptic facilitation in Syt7KO mouse^27^. The longer isoform includes extra amino acids of unknown function in the spacer domain localized between the transmembrane region and the C2A domain. We observed a dramatic decrease in the expression of the 46 kDa splice variant of Syt7 (**Fig. 3d,e**) in agreement with decreased staining in brain slices (**Fig. 3a** and **Supplementary Fig. 3a**).

**Figure 3.**
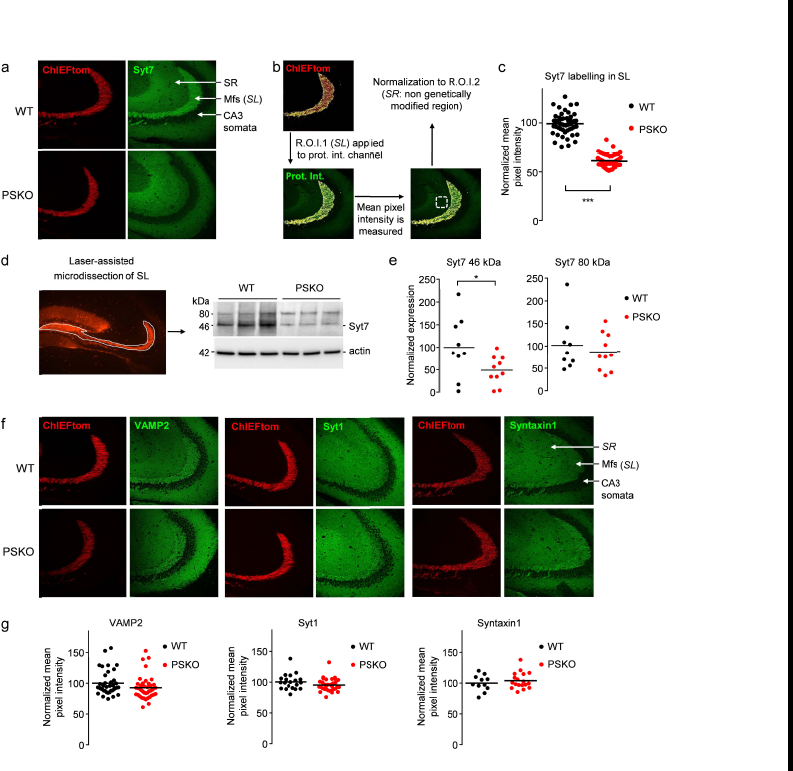
PS specifically supports the expression of Syt7. (**a**) Confocal images of the *stratum lucidum* (SL) where the Mfs can be distinguished in the red channel (ChIEFtom) and stainings of Syt7 depicted in green. The Syt7 staining in PSKO Mfs was negligible in comparison to WT. (**b**) To quantify the intensity of the stainings a mask corresponding to the *SL* was created in the red channel. This first region of interest (R.O.I.) was applied to the green channel were the mean pixel intensity of the protein of interest was measured only in the *SL.* To normalize the results, the mean pixel intensity was also measured in a second R.O.I.: the *stratum radiatum* (SR). The staining intensities in the *SL* were normalized to the intensities in the SR. (**c**) Scatter plots of the mean pixel intensity in the *SL* normalized to the mean pixel intensity in the *SR.* A reduction of 40% of intensity is observed while approximately half of the Mfs are projected from PSKO GCs. n=53 slices from 9 mice in WT, n=50 slices from 8 mice in PSKO, *p=0.0004.* (**d**) *Left*, confocal image of the hippocampus depicting the area dissected with a laser to prepare protein lysates. *Right*, western blot from these lysates showing a decreased expression of Syt7. (**e**) Quantitative analysis of western blot in “c”. A 50% decrease in the expression of the 46kDa band of Syt7 was detected. (**f**) Confocal images of the *stratum lucidum (SL)* where the Mfs can be distinguished in the red channel (ChIEFtom) and stainings of VAMP2, Syt1 or Syntaxin1A are seen in green. (**g**) Scatter plots of the mean pixel intensity in the *SL* normalized to the mean pixel intensity in the *SR* measured in images as in “e”. Staining patterns and intensities do not differ between genotypes. For VAMP2, n=38 slices from 8 mice in WT, n=41 slices from 9 mice in PSKO. For Syt1, n=20 slices from 5 mice in WT, n=33 slices from 7 mice in PSKO. For Syntaxin1, n=11 slices from 4 mice in WT, n=18 slices from 7 mice in PSKO.

We then tested whether the functional impairment of SV release at PSKO Mf-CA3 synapses could result from fewer SVs available or from a limited amount of molecules necessary for SV release. To test these possibilities, we labeled brain slices against VAMP2, the most abundant SV protein; against synaptotagmin 1 (Syt1), the Ca^2+^ sensor necessary for synchronous SV release; against syntaxin1, a SNARE complex protein integrated at the plasma membrane necessary for the release of SV (**Fig. 3f);** and against Bassoon (**Supplementary Fig. 2g**). We measured the mean pixel intensity of staining in the *SL* and did not observe any difference between genotypes for these presynaptic markers (**Fig. 3g**). Thus, the deficit in Syt7 expression appears rather specific among proteins of the presynaptic machinery.

### Impaired facilitation and SV replenishment in PSKO Mfs is caused by decreased Syt7 expression

We thus postulated that impaired facilitation and SV replenishment at PSKO Mf-CA3 synapses are the functional consequences of decreased Syt7 expression. To address this hypothesis, we tested whether the molecular re-expression of Syt7 in PSKO Mfs was sufficient to recover the functional impairment in synaptic plasticity and SV replenishment rate. We used a strategy involving an injection of PSfl mice with two viral vectors: a first virus allows expression of Cre in GCs leading to deletion of PS and to expression of Cre-dependent ChIEFtom and of Syt7 which is brought by a second virus (**Fig. 4a**). While GCs infected by only one virus do not respond to light, double-infected GCs lack PS, yet express Syt7 and can be controlled by light. Under this condition, in PSKO, Syt7 was correctly re-expressed in Mf axons (**Fig. 4b**). The EPSCs evoked by light stimulation in GC dendrites displayed basal synaptic properties similar to WT or to PSKO Mf-CA3 synapses (**Supplementary Fig. 4a**). However, presynaptic facilitation in response to high frequency trains of stimulation (**Fig. 4c,d**) and the SV replenishment rate (**Fig. 4e,f**) recovered to levels not significantly different from WT levels. Hence, normal synaptic properties can be rescued in PSKO Mf-CA3 synapses by re-expression of Syt7, demonstrating that the decreased expression of Syt7 is responsible for the PSKO functional phenotype.

**Figure 4.**
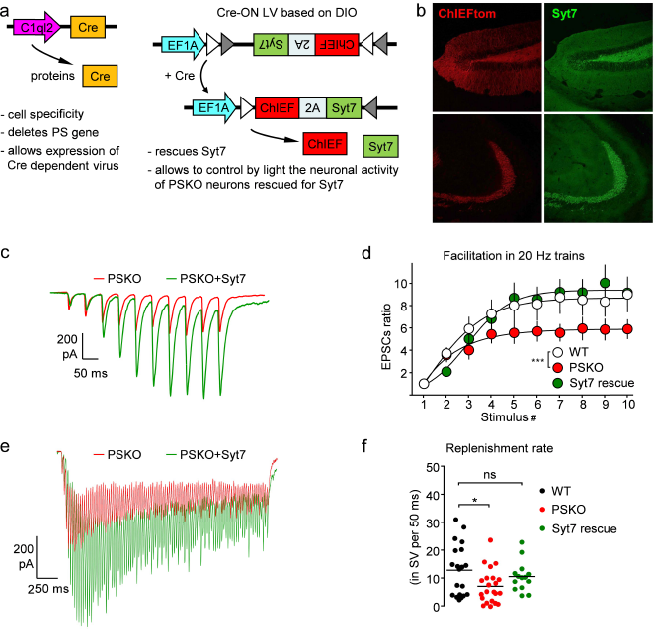
Impaired facilitation and SV replenishment in PSKO Mfs is caused by decreased Syt7 expression. (**a**) Schemes of the two lentiviruses (LV) used to rescue Syt7 expression in PSKO GCs controlled by light. C1ql2 promoter restricts the expression of Cre to GCs where the floxed PS gene is deleted. The Cre also allows the Cre-ON LV to express Syt7 and ChIEF. (**b**) Confocal images showing the rescued expression of Syt7 in the hilus (up) and in the *stratum lucidum* (down) using the strategy described in “a”. (**c**) Representative traces of EPSCs evoked by ten repetitive light-pulses at 20 Hz. ( **d**) Analysis of presynaptic facilitation in 20 Hz trains. Facilitation is represented by the amplitude of the EPSCs normalized to the amplitude of the first EPSC. Facilitation, impaired in absence of presynaptic PS, recovers to WT level by re-expression of Syt7. (**e**) Representative EPSCs evoked by 100 stimuli at 40 Hz. The amplitude of EPSCs is rescued by Syt7 re-expression. (**f**) Scatter plots with means of replenishment rates calculated per neuron are represented by genotype. The replenishment rate impaired by the absence of PS is rescued by re-expression of Syt7.

### γ-secretase activity regulates Syt7 protein levels

We then asked whether the loss of γ-secretase proteolytic activity, normally mediated by PS, was involved in the decreased expression of Syt7. We investigated this possibility by treating rat neuronal cultures with a γ-secretase inhibitor (GSI: L685,458 at 10 μM) for four days (from 10 to 14 DIV) and preparing protein lysates for western-blot (WB) analysis. The Syt7 antibody labelled two bands at 46 and 80 kDa (**Fig. 5a),** as in the microdissected brain samples. Interestingly, the intensity of the lowest band was decreased by half in conditions of GSI treatment (**Fig. 5a,c),** indicating that blocking γ-secretase proteolytic activity downregulates Syt7 expression. In contrast, the expression levels of VAMP2, Syt1 and Syntaxin1 (**Fig. 5b,d**) were not modified by GSI treatment. Hence, the regulation of Syt7 by γ-secretase activity is specific among the presynaptic markers tested.

**Figure 5.**
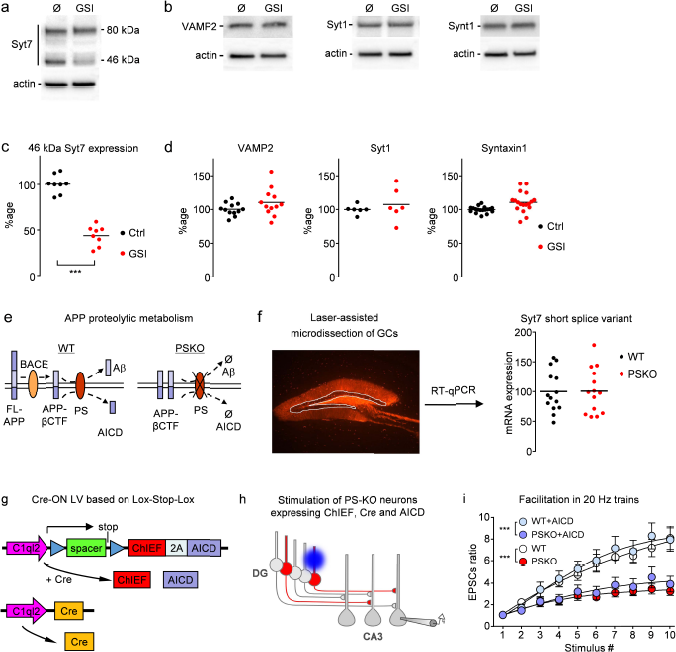
γ-secretase activity regulates Syt7 protein levels. (**a-b**) Western-blots of protein samples from neuronal cultures treated or not with 10 |jM GSI from 10 to 14DIV. The 46kDa Syt7 band decreases dramatically when γ-secretase is inhibited. γ-secre tase inhibition does not impair expression of VAMP2, Syt1 and Syntaxin1A. (**c-d**) Scatter plots of the western-blot quantification of samples from neuronal culture treated or not with GSI. Syt7 expression is decreased by half in treated condition as seen in “a” while the expression of VAMP2, Syt1 and Syntaxin1A is stable under GSI condition as seen in “b”. (**e**) Schemes of APP proteolytic metabolism in WT and PSKO conditions. Full-length (FL) APP is first cleaved by BACE producing βCTF, a transmembrane stub, further degraded by PS (**y**-secretase) into Ap and APP IntraCellular Domain (AICD). In absence of PS, βCTF cannot be catabolized and thus accumulates. (**f**) *Left*, confocal image of the hippocampus depicting the GCs layer dissected with a laser to prepare RNA samples. *Right*, RT-qPCR analysis of the samples indicates that mRNA level of Syt7 is not impaired in absence of PS. (**g**) Schemes of the two lentiviruses (LV) used to rescue AICD expression in PSKO GCs controlled by light. C1ql2 promoter restricts the expression of Cre to GCs where the PS floxed gene is deleted. The Cre also removes the Lox-Stop-Lox sequence allowing the Cre-ON LV to express AICD and ChIEF. (**h**) Strategy to selectively study Mf/CA3 synapses that are presynaptically PSKO, yet express AICD. In PS *“floxed”* conditional knock-out mice, double-infected GCs do not express PS (Cre deletes PSfl gene), yet express AICD (Cre dependently) and ChIEF (bicistronic construct) allowing the control of their activity by light. GCs infected by a single virus do not respond to light. (**i**) Facilitation of EPSCs amplitude measured along trains of 10 stimuli at 20 Hz in WT or PSKO expressing AICD. The impaired facilitation of PSKO is not rescued by expression of AICD. Two-way ANOVA, n=8 in WT, 11 in PSKO, *p=0.017*.

This involvement of γ-secretase activity indicates a possible role for γ-secretase substrates expressed at the presynaptic site. APP, the main proteolytic substrate of γ-secretase is first processed by β-secretase (BACE1) producing the βCterminal fragment of APP (APP-βCTF), a transmembrane stub further cleaved by the intramembrane proteolytic activity of γ-secretase releasing Aβ peptides and the APP-intracellular domain (AICD) (**Fig. 5e**). We hypothesized that the PSKO phenotype, including the decreased expression of Syt7 results from the absence of AICD because this γ-secretase product has been reported to regulate transcription^31^. We performed laser capture microdissection of the GC layer to prepare RNA samples used for qRT-PCR of Syt7 mRNA, and did not observe any change between the two genotypes (**Fig. 5f**). We further addressed the possible involvement of AICD in presynaptic mechanisms by testing whether the molecular rescue of AICD in PSKO GCs could recover the functional phenotype. Once again we used a strategy based on the combination of a Cre-expressing lentivirus with a Cre-ON lentivirus (**Fig. 5g**). This allowed the optical control of activity of GCs which were PSKO but still expressed AICD (**Fig. 5h**). In this condition, we did not observe any recovery of presynaptic facilitation (**Fig. 5i**) while basal transmission was not affected (**Supplementary Fig. 5a,b**). The lack of evidence for an impairment at the transcriptional level indicates that Syt7 protein is dysregulated post-transcriptionally.

### Syt7 downregulation results from the accumulation of the γ-secretase substrate APP-βCTF

In the absence of PS and its γ-secretase proteolytic activity, unprocessed APP-βCTF substrates may accumulate. We studied the expression level of different fragments of APP by staining brain slices with antibodies directed against either the Nterminal or Cterminal epitopes of APP. In control conditions, we observed that the *SL* was moderately stained with the Cterminal antibody relatively to the CA3 somata and barely stained with the Nterminal antibody (**Fig. 6a**). Interestingly, in conditions where PS is selectively suppressed in Mf axons, we observed a striking increase of APP staining with the Cterminal but not with the Nterminal antibody indicating that an APP fragment lacking the Nterminal epitope accumulates in PSKO axons (**Fig. 6a,b**). Using protein samples obtained by laser microdissection of PSKO Mfs, we distinctly identified APP-βCTF as the APP fragment accumulating in Mf axons (**Fig. 6c,d**). Thus, in PSKO Mfs, APP is first cleaved by BACE1 which is abundantly expressed in Mfs (ref. ^32^ and **Fig. 6e),** and produces APP-βCTF which accumulates in Mf terminals demonstrating that it can neither be retrogradely transported nor be catabolized by enzymes other than PS. We induced an accumulation of APP-βCTF in neuronal cultures similar to the one observed in Mfs by blocking PS function with a γ-secretase inhibitor (GSI). Interestingly, we observed an inverse correlation between Syt7 expression level and APP-βCTF which progressively accumulates in protein lysates from cultures treated from 6 hours to 9 days with GSI (**Fig. 6f**) suggesting that accumulation of APP-βCTF prevents the normal expression of Syt7.

**Figure 6.**
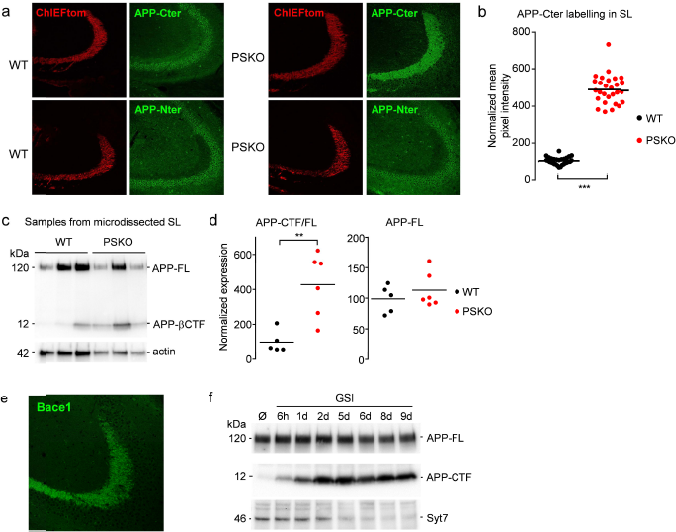
γ-secretase proteolytic activity regulates the axonal expression of APP-βCTF. (**a**) *Left*, confocal images of the Mfs tracts in WT distinguished in the red channel (ChIEFtom) and of stainings of APP Cterminal with Y188 antibody (upper panel) and Nterminal with JRD32 antibody (lower panel). Mfs are labelled with the Cterminal but barely with the Nterminal antibody. *Right*, in PSKO, Mfs are strongly labelled with the Cterminal but not with the Nterminal antibody. (**b**) Samples prepared as in “Fig.3c” show a striking increase in APP-βCTF levels in dissected Mfs detected with Y188 antibody against APP-Cter. (**c**) Quantitative analysis of western blot in “B”. A 350% increase in the expression of APP-βCTF was detected. (**d**) Immunolabelling of β-secretase BACE1 in brain sections. BACE1 expression is prominent in the *SL* where Mfs contact CA3 pyramidal cells. (**e**) Western-blot of samples prepared from neuronal cultures treated with GSI from 6 hours to 9 days *in vitro.* APP-CTF accumulates while Syt7 drops.

Amongst other γ-secretase substrates identified so far only a few of them are known to be expressed at the presynaptic site and could potentially play a role in the presynaptic phenotype observed^33,34^. We identified N-cadherin and Neurexin1 as additional candidate substrates that may accumulate in presynaptic Mf terminals upon deletion of PS. We studied the accumulation of the transmembrane stubs of these proteins produced after ectodomain shedding in neuronal cultures treated with GSI (**Supplementary Fig. 6a-d**). We also stained hippocampal slices using antibodies against the Cterminal fragments of these two proteins. We observed, as already reported, that Ncad-CTF1 accumulates in neuronal cultures^35^ but not in PSKO Mf terminals. The Cterminal stub of Neurexin does not accumulate in cultured neurons treated with GSI nor in PSKO Mfs (**Supplementary Fig. 6a-d**).

To address whether there was a causal relation between APP-βCTF accumulation and Syt7 downregulation, we crossed the PS1fl mouse line with an APPfl mouse line^36^. In the resulting APPfl/PS1fl mouse line, the expression of Cre removes both PS1 and APP in the same cells; thus even though the enzyme is removed, no substrate (APP-βCTF) can accumulate (**Fig. 7a**). We compared the expression level of Syt7 in APPfl/PS1fl to PS1fl and WT (**Fig. 7b,c**). In PS1KO Mfs from PS1fl mice, the level of Syt7 expression was significantly decreased compared to WT. However, the extent of this reduction was less pronounced than the reduction observed in PSKO Mfs from PS1fl/PS2KO mouse. This is in agreement with the milder impairment of presynaptic facilitation observed in PS1KO compared to PSKO (removal of both PS1 and PS2) conditions (**Supplementary Fig. 2a**) and suggests possible compensatory changes in PS2 expression in a PS1KO background, as observed in neuronal cultures^35^. In Mfs expressing neither PS1 nor APP, the expression level of Syt7 was comparable to WT. Thus the absence of APP prevents the decreased expression of Syt7 in Mfs, demonstrating a causal relationship between APP processing and downregulation of Syt7. While the level of Syt7 differs between PSKO and APPKO/PS1KO conditions, γ-secretase products of APP processing (A**p,** AICD) cannot be produced in both conditions, indicating that the Syt7 deficit does not result from the absence of Aβ or AICD. In contrast, accumulation of APP Cterminal stubs can occur in PSKO but not in APPKO/PS1KO cells establishing APP-βCTF as the causative APP fragment involved in Syt7 regulation.

**Figure 7.**
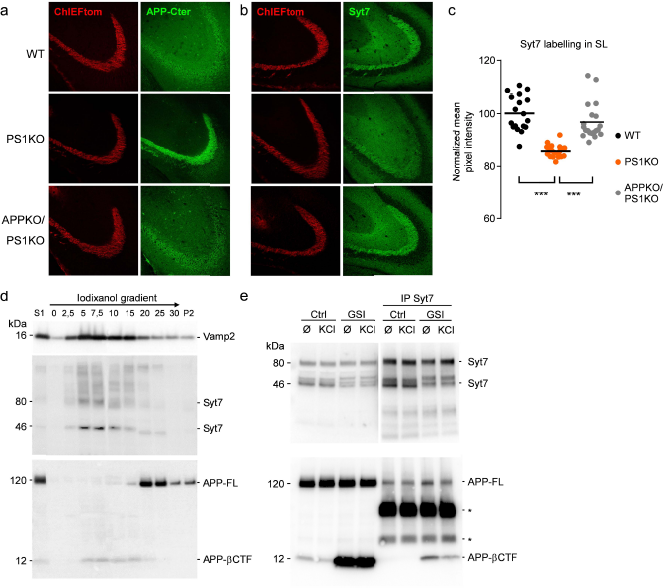
PS regulation of Syt7 expression depends on APP-βCTF, a new interacting partner of Syt7. (**a**) Confocal images of ChIEFtom (red channel) and APP-Cterminal (detected with C1/6.1 antibody, green channel) in the Mf tracts in WT, PS1KO and PS1KO/APPKO conditions. Mfs are faintly labelled with the APP antibody in WT and barely in APPKO/PS1KO. In contrast, PS1KO Mfs display strong labelling of APP. (**b**) Confocal images as in *(a)* displaying Syt7 labelling. Mfs are strongly labelled in WT and in APPKO/PS1KO but faintly in PS1KO. (**c**) Scatter plots of the mean pixel intensity in the *SL* normalized to the mean pixel intensity in the *stratum radiatum* report a reduction of 18%. In this condition, 50% of the GCs are infected by a Cre-expressing LV thus approximately 50% of the Mfs are projected from non-infected (PS expressing) GCs. (**d**) Western-blot of cellular fractions from cultured neurons separated on an iodixanol gradient. VAMP2, Syt7 and APP-βCTF are enriched in light fractions while APP-FL is present in heavier fractions. (**e**) Samples prepared from cultured neurons treated or not with GSI from 10 to 14 DIV and with or without KCl (30 mM) for 15′ before lysis. In the inputs, detection of APP Cterminal with the Y188 antibody indicates that GSI induces β CTF accumulation. IP of Syt7 pulled-down APP-FL and β CTF. Asterisks indicate light and heavy chains of the immuno-precipitating antibody. Syt7**/p**CTF interaction is decreased by KCl treatment suggesting that it is regulated by neuronal activity.

We further investigated the relationship between APP-βCTF and Syt7 by performing cell fractionation on an iodixanol gradient. We observed that Syt7 was enriched in the lighter fractions from neuronal culture extracts characterized by high expression of VAMP2 (**Fig. 7d**). Interestingly, APP-βCTF was detected in the same fractions while full-length APP was expressed in the more heavy fractions (**Fig. 7d**). This suggests a possible interaction between APP-βCTF and Syt7 in the lighter cellular compartments of the neurons. We thus studied a possible interaction between Syt7 and the APP fragments. Immunoprecipitation of Syt7 from protein lysates of neuronal cultures allowed co-precipitating APP indicating that the two proteins interact (**Fig. 7e**). When neurons were treated with GSI, APP-βCTF dramatically accumulated and was massively co-precipitated with Syt7 (**Fig. 7e**). Moreover, when the neuronal cultures were treated with KCl to depolarize the neurons and induce action potential firing, the interaction between Syt7 and APP-βCTF decreased, indicating that the interaction was dynamically regulated by neuronal activity and possibly by SV release (**Fig. 7e**).

## Discussion

Understanding the physiological function of AD-related genes is fundamental because dysregulation of their normal function likely contributes to AD pathogenesis. Because APP and PS are abundantly expressed in presynaptic compartments where they interact with proteins of the release machinery ii is essential to understand their physiological role in presynaptic mechanisms. To address this, we have developed versatile optogenetic tools that help circumvent difficulties inherent to studies of presynaptic mechanisms with classical electrophysiological methods. Indeed, following genetic manipulations in a subpopulation of neurons, stimulation with an extracellular electrode activates both the axons from genetically manipulated cells and those from non-modified cells. We cloned a new minimal promoter of the C1ql2 gene and used it in a lentiviral vector to target expression of ChIEF in DG together with a second transgene, e.g Cre recombinase. C1ql2-driven expression of ChIEF and Cre allowed the combination of region specific activation with gene deletion in the DG of PS-conditional KO mice. Using this approach, we examined the effects of the genetic deletion of PS in presynaptic GCs on the morpho-functional properties of Mf-CA3 synapses.

### PS/γ-secretase regulates presynaptic mechanisms by supporting the expression of Syt7

We discovered that PS controls the expression of the Ca^2+^ sensor Syt7 presynaptically. Indeed, in the absence of presynaptic PS, the axonal expression of Syt7 dramatically drops. Interestingly, this phenomenon seems rather specific for Syt7 as four other presynaptic markers tested were not impaired by the absence of PS, two being SV proteins (VAMP2, Syt1) and two being active zone proteins (Bassoon, Syntaxin1A). Moreover, the absence of PS does not impact on the morphology of presynaptic terminals indicating, all together, that the functional phenotype is not caused by degenerative processes within presynaptic terminals.

The reduced expression of Syt7 in PSKO Mfs impairs two essential presynaptic functions: presynaptic facilitation and replenishment of release-competent vesicles. Indeed, Syt7 has recently been shown to be essential for presynaptic facilitation^27^ at several central synapses. Its role is especially noteworthy at Mf-CA3 synapses where the large presynaptic train facilitation is nearly abolished in Syt7KO^27^. Thus, the impaired short-term plasticity observed at PSKO Mf-CA3 synapses is consistent with the decreased expression of Syt7 in Mf terminals. Additionally, Syt7 is necessary for the replenishment of release competent SVs^28^. We demonstrated that re-expression of Syt7 in PSKO Mf-CA3 synapses rescued presynaptic function, providing a causal link between the drop of Syt7 expression and the functional phenotype. Of note, quantal and basal transmission were unchanged in agreement with the observation that Syt7 does not determine the initial probability of release^27,28^.

Next, we investigated whether this novel synaptic function of PS depends on its γ-secretase proteolytic activity. In neuronal cultures, we observed that inhibiting γ-secretase activity pharmacologically decreased Syt7 expression within a few days indicating a role for γ-secretase. The investigation of molecular mechanisms involving γ-secretase is complex due to its diverse functions and molecular targets^8^. APP is a major substrate of PS and is highly enriched in synapses, and particularly at a presynaptic level^12^. In the absence of PS or when γ-secretase activity is inhibited, the substrate of PS, APP-βCTF, cannot be processed into Aβ and AICD. Thus, in the PSKO condition, only the absence and not the excess of Aβ and AICD could potentially explain the PSKO phenotype. Because AICD has been proposed to act as a transcription factor^31^, the decline in Syt7 level could result from a deficit in AICD. However, the analysis of Syt7 mRNA and rescue experiments of AICD expression in PSKO GCs show that the absence of AICD does not play a role in the regulation of Syt7 expression and function.

### PS/γ-secretase regulates presynaptic facilitation by controlling the level of APP-βCTF

We observed that the substrate of PS, APP-βCTF, accumulates in Mf axons in absence of PS, or in cultured neurons when γ-secretase is pharmacologically inhibited. This fragment of APP is involved in the deficit in Syt7 expression observed in PSKO axons since deletion of APP in APPKO/PS1KO prevents the drop of Syt7 expression. Moreover, there was an inverse relationship between APP-βCTF and Syt7 expression in neuronal cultures. Finally we show that Syt7 is a new interacting partner of APP-βCTF. The mechanism linking the accumulation of APP-βCTF to downregulation of Syt7 will need further investigation. A possible hypothesis is that βCTF sequesters Syt7 in compartments preventing its functional roles and causing its degradation. This interpretation is supported by the well documented function of PS and APP in axonal transport^37–40^ and by the observation that the absence of PS impairs the cell surface expression and the shuttling of several proteins between the cell interior and the cell surface^41,42^. Syt7, as a protein expressed both in SVs and at the plasma membrane^43,44^ could be sequestered in the wrong compartment by accumulation of APP-βCTF. Our results are based on the pharmacological inhibition of γ-secretase activity or genetic invalidation of PS. They suggest that in physiological conditions, presynaptic γ-secretase activity regulates the abundance of Syt7 a key presynaptic protein involved in presynaptic plasticity which interacts with APP-βCTF. It will be interesting to investigate the physiological conditions under which γ-secretase activity is regulated, hence modulating the amount of APP-βCTF and consequently of Syt7.

Although our results mainly deal with the investigation of the physiological role of PS, they may also be of importance for the physiopathology of AD. FAD mutations of PS1 also cause APP-βCTF accumulation due to decreased enzymatic activity of γ-secretase, limiting the catabolism of APP-βCTF^45^. Interestingly, genetic evidence indicates a relationship between the level of APP-βCTF catabolized by γ-secretase and AD. Indeed, APP-βCTF is highly produced when the APP gene contains mutations causing the early onset form of AD (the Swedish mutations K595N/M596L^2^) and is produced to a much lower extent when APP contains a mutation (A598T) protecting from AD^46^. Overall our data highlight the notion that presynaptic dysfunction may play a role in AD in relation to APP processing by PS.

## AUTHOR CONTRIBUTIONS

G.B., T.J.S., J.R-M and F.B. conducted the experiments, G.B., U.M. and C.M. designed the experiments and wrote the manuscript.

## ACKNOWLEDGEMENTS

We thank John Lin from UCSD-HHMI for his generous gift of ChIEFtdTomato plasmids. Lentiviruses were produced at the TransBioMed facility of the Bordeaux University. Cellular imaging was performed at the Bordeaux Imaging Center, a service unit of the CNRS-INSERM and Bordeaux University, member of the national infrastructure France BioImaging. Microdissections assisted by laser capture and RT-qPCR were performed at the facilities of the Neurocentre Magendie. Gael Barthet is supported by a young researcher position from the “Fondation Plan Alzheimer” and by a “return-post-doc” fellowship form the French national agency for research (ANR). This project has received funding from the European Union’s Horizon 2020 research and innovation programme under grant agreement No 676144. Ulrike Müller is supported by a grant from the Deutsche Forschungsgemeinschaft (MU 1457/14-1). The authors declare no competing financial interests. We are grateful to Christophe Blanchet, Thierry Amédée and Ruth Betterton for their careful comments on the manuscript.

## METHODS

### Ethical approval

Animal anesthesia and euthanasia procedures were carried out in accordance with the Animal Protection Association of ethical standards and the French legislation concerning animal experimentation (authorization #A33 12 061).

### Animals

The animals used in this study were PS1floxed #007605, PS2-KO #005617 mice obtained from Jackson Laboratory (Bar Harbor, ME, USA) and C57BL6/J Wild Type (WT). In the PS1floxed mice, loxP sites are present on either side of exon 7 of the targeted gene and allow deletion of this exon by Cre-mediated recombination. In the PS2 mouse line, a targeting vector was used to disrupt exon 5 and introduce a frame shift between exons 4 and 6. We developed a PS1fl/PS2KO mouse line by crossing the PS1floxed and the PS2KO mice together. Mice were genotyped using the following primers: PS1fl-F: CAGGCACACTCACCCACAGA, PS1fl-R: GAAAATCATATCCCCTACACTA; PS2WT-F: GAGAGAAGGCACCAGGATAGGTT, PS2WT-R: AGTGTCCTGGAGCAGCAGATAC and PS2Mut-F: CGAAGGAGCAAAGCTGCTATTGG, PS2Mut-R: CTGGGTCTCCAGCAGTCTTC. We developed an APPfl/PS1KO mouse line by crossing the APPfloxed^36^ and the PS1floxed mice together. APP gene was genotyped using the following primers: 1: AACGCAGGGAGGAGTCAGGG, 2: TGCATGTCAGTCTAATGGAGGC, 3: ATCTGCC CTTATCCAGTGAAATGAACC.

### Cloning of C1ql2 minimal promoter and generation of lentiviral vectors

The lentiviruses were produced by the viral vectors facility of Bordeaux “TransBioMed”. The lentiviruses developed here are based on the pRRLsin-PGK-MCS-WPRE plasmid where PGK promoter has been exchanged with the C1ql2 minimal promoter. This sequence was cloned from mouse genomic DNA using the following primers EcoRV/C1ql2-F: ATATATGATATCagcacccacatagcagc; BamH1/C1ql2-R: ATATATGGATCCgctctggaactgat ctg. The 1.3kb sequence was then inserted between EcoRV and BamH1 restriction sites of the lentiviral backbone. After the promoter, the following cDNA sequences were inserted in the 5’ to 3’ order: ChIEFtomato, 2A peptide from porcine teschovirus-1^47^, finally the Cre recombinase or the eGFP. A Cre-dependent construct was created to allow conditional expression of ChIEFtom and AICD. Behind C1ql2 promoter, a Lox-Stop-Lox cassette was inserted followed by the cDNA of ChIEFtom, 2A peptide and AICD. A second Cre-ON construct was created based on a double-inverted open-reading frame (DIO) strategy. Briefly, a cassette including ChIEFtom, 2A peptide and Syt7 was placed in an inverted manner between two doublets of lox sequences (loxP and lox 2722). For 3D reconstruction of Mf boutons, a lentiviral vector expressing myristoylated YFP (mYFP) was used (generously provided by C. Lois, California Institute of Technology, Pasadena, CA) described in^26^.

### Viral gene transfer and stereotaxic delivery

Mice (P22-P30) were anesthetized by isoflurane inhalation and injected with buprenorphine to prevent post-surgery pain. 1 μl of viral solution (5.10^6^ particles) was injected using a micropump and syringe (Nanofil WPI) at the rate of 100 nl/min in the DG (Y: 2.2 mm from lambda; X: ± 2.0 mm from sagittal suture; Z: 2.0 mm from the skull) to infect GCs. This site targets a rather dorsal part of the hippocampus but the spreading of viruses allowed the infection of a large part of the hippocampus (radius of roughly 1 mm) to produce numerous slices for IHC or electrophysiology. Experiments were performed at least 3 weeks post-injection.

### Slice preparation

Mice were anesthetized using ketamine/xylasine mix (100 mg/kg /10 mg/kg ; i.p.) and perfused transcardially with ice cold artificial cerebrospinal fluid (aCSF) for 40-60 s. The brain was quickly removed from the skull and chilled in ice-cold aCSF containing the following (in mM): 120 NaCl, 26 NaHCO_3_, 2.5 KCl, 1.25 NaH_2_PO_4_, 2 CaCl_2_, 1 MgCl_2_, 16.5 glucose, 2.8 pyruvic acid, 0.5 ascorbic acid, pH 7.4 adjusted by saturating with carbogen (95% 02 and 5% CO2), and an osmolarity of 305 mOsm. Parasagittal slices (300 μm) were cut from brain hemispheres using a vibratome (VT 1200S, Leica Microsystems, Nussloch, Germany). Slices were then transferred to aCSF for 20 minutes at 33°C and thereafter maintained at room temperature until required.

### Electrophysiological recordings

A slice was transferred to a recording chamber where it was submerged and continuously perfused with oxygenated (95% 02, 5% CO2) aCSF at RT. Whole-cell patch clamp recordings were performed on neurons identified with a differential interference contrast microscope (Eclipse FN-1, Nikon, Champigny sur Marne, France) equipped with a camera (CoolSNAP EZ, Ropper, Evry, France). Voltage-clamp recordings were performed on CA3 pyramidal neurons using patch electrode (4±0.5 MΩ) filled with an internal solution containing the following (in mM): 125 CsCH_3_SO_3_, 2 MgCl_2_, 4 NaCl, 5 phospho-creatine, 4 Na_2_ATP, 10 EGTA, 10 HEPES pH 7.3 adjusted with CsOH and an osmolarity of 300 mOsm. Bicuculline (10 μM) was added to the bath to inhibit GABA-A receptors. Mf-CA3 excitatory post synaptic currents (EPSCs) were evoked by light-pulses of 1 ms over GCs dendrites in the *stratum moleculare*. The baseline stimulation frequency for all experiments was 0.1 Hz. Current-clamp recordings were performed on GCs or CA3 cells patched with an internal solution containing the following (in mM): 115 KC_6_H_11_O_7_, 10 KCl, 0.02 CaCl_2_, 4 MgATP, 0.3NaGTP, 15 phospho-creatine, 0.2 EGTA, 10 HEPES pH 7.3 adjusted with KOH and an osmolarity of 300 mOsm. Spike transmission experiments were performed in current clamp mode while using the “low frequency voltage clamp” function of EPC10 amplifier to keep the membrane potential of the recorded CA3 neurons constant at -70 mV between recording sweeps. Signals were amplified using a HEKA EPC10 amplifier (Lambrecht, Germany), filtered at 3.3 kHz and digitized at 10 kHz via PatchMaster software (Lambrecht, Germany). Data were analyzed using IGOR PRO 6.3 (Wavemetrics, Lake Oswego, OR).

### Analysis of quantal transmission

The characteristics of quantal neurotransmission at Mf/CA3 were determined by recording EPSCs in CA3 neurons while stimulating with light-pulses at 1 Hz in presence of 32 μM of cadmium. We used the following model to estimate the probability of release at individual releasing site (active zone) and we estimated the contamination of mono-quantal release by bi-quantal release in the experiments in presence of 32 μM Cadmium. In this condition, the failure rate is of 88%, hence the probability of release is 12%. This total probability is the sum of the probability of release at individual release sites (we considered 20 sites/Mf bouton here) and possibly of concomitant release from two sites (concomitant release from more than two sites are negligible and not considered here). For the simplicity of the demonstration, we postulate the release sites to be equivalent.

Thus, P(tot) = 0.12 = P(s1) + P(s2) + … + P(s20) + P(s1+s2) + P(s1+s3) +…+ P(s19+s20) = 20P(s) + 190 P(s+s)

The equation of second order is: 190 P(s+s) + 20P(s) - 0.12 = 0 and its solutions are: P(s)1 = - 0.11095 and P(s)2 = + 0.00569

Thus, P(s+s) = (0.12- 20*0.00569)/190 = 0.0000316. In our recordings, the ratio of release from two sites compared to one site is: (190*0.0000316)/ (20*0.0057) = 0.0526 i.e. 5.26%. In our condition (32 μM of Cadmium) for the analysis of quantal release, 5% of the EPSCs were possibly coming for the concomitant release from two sites.

### Immunostaining

WT and PSKO mice were anesthetized with intraperitoneal administration of pentobarbital (50 mg/kg body weight) and were perfused intracardiacally with 0.9% NaCl for 1 min followed by 4% paraformaldehyde in 0.1 mM PBS for 3 min for imaging experiments. Brain were removed and postfixed in 4% PFA for 8 h. Coronal sections of the hippocampus (50 μm) were cut on a vibratome (Leica VT1000S). Sections were permeabilized in 0.3% TritonX100 for 2h at room temperature. Then sections were incubated with primary antibody in PBS-tween for 48 h at 4°C (APP N-terminal JRD32 mouse antibody is a non-commercial antibody directed against the N-terminal APP-E1 domain^48^, APP C-terminal Y188 rabbit antibody Abcam #32136 or APP C-terminal C1/6.1 mouse antibody BioLegend, BACE1 rabbit antibody D10E5 Cell Signaling #5606; syntaxin1 rabbit antibody Alomone # ANR-002; from Synaptic Systems: bassoon rabbit antibody #141013, synaptoporin rabbit antibody #102002, synaptotagmin1 mouse antibody #105011, synaptotagmin7 rabbit antibody #105173, synaptobrevin2 mouse antibody #104211). After washes in PBS-T, sections were incubated with secondary antibodies for 3h (coupled to either Alexa 488 or 555 or anti-rabbit Abberior STAR 635P). Slices were incubated during the penultimate wash in DAPI (300 nM) for 10’ before a last PBS wash and mounted in Mowiol based solution. For low magnification acquisition, an up-right widefield microscope (Zeiss AxioPhot2) was used with a 2.5X objective. The camera was a CoolSnap HQ2 (Photometrics, Tucson, USA) controlled by MetaMorph.

### Dual hemisphere acquisition after unilateral conditional deletion of PS gene

WT or PS1fl/PS2KO mice were injected with lentivirus encoding ChIEFtom2A-Cre on the right hemisphere and ChIEFtom2A-GFP on the left hemisphere using stereotaxic surgery. After fixation and brain slicing, whole brain sections, fluorescently labelled against synaptotagmin1, synaptotagmin7 or synaptobrevin2 were acquired using the digital slide scanner NanoZoomer 2.0HT (Hamamatsu).

### Confocal laser-scanning fluorescence microscopy of mossy fiber tract and image analysis

Confocal acquisition were performed using a Leica SP8 White Light Laser 2 on an inverted stand DMI6000 (Leica Microsystems, Mannheim, Germany) and with a 20X (NA 0.75) objective. Images of focal planes in the region of the *stratum lucidum* were acquired in the red (ChIEFtomato) and far-red (synaptotagmin1, synaptotagmin7, synaptobrevin2, syntaxin1) channels. The settings (laser power, gain, offset) were fixed between genotypes. Images were analyzed using ImageJ. A region of interest (ROI) restricted to the Mf tract was defined in the red channel using the default mode threshold of ImageJ followed by the creation of a selection corresponding to this ROI. Then average pixel intensity was measured in the same ROI but in the far-red channel. The mean pixel intensity of a second area (ROI2) in the *stratum radiatum* (SR), an adjacent region not genetically manipulated was measured. A ROI/ROI2 ratio was performed slice per slice to normalize the data (fig. 3B). Between 40 and 50 measurements from 6 to 8 mice were acquired per genotype and per presynaptic proteins studied.

### Mossy fiber boutons analysis after 3D-stack acquisition

Confocal acquisition was performed using a Leica SP8 described above and with a 63X (NA 1.40) objectives. Images of the Mf boutons expressing myristoylated YFP in the CA3b-c region of hippocampus were captured with an interval of 0.2 μm in z-axis and a pixel size of 70 μm. 200 Mf boutons from five animals per genotype were analyzed. High-resolution 3D stacks were generated using Imaris 7.1.1 software (Bitplane) to characterize and analyze presynaptic terminals. Illustrations correspond to 3D reconstructions smoothened by a median filter.

### STED microscopy and image analysis

STED acquisitions were performed on the same microscope using a 775 nm depletion laser and a 100X (NA 1.40) objective. Images of the Mf bouton were first acquired on the yellow channel (myristoylated YFP) to obtain a mask of the Mf bouton. An intensity threshold was applied to define the region contour of the Mf bouton. Then, a STED image of bassoon puncta was acquired in far-red with a pixel size of 20 nm. To count the number of bassoon clusters per Mf bouton and to measure their surface, images were treated by a segmentation program allowing clusters to be detected as single objects, which were then automatically counted within the contour of Mf boutons. Analyses were performed with Image J software (National Institutes of Health, New York, USA).

### Laser assisted microdissection and RTqPCR

Laser capture microdissection (LCM) was performed using a MicroBeam microdissection system Version 4.8 equipped with a P.A.L.M. RoboSoftware (P.A.L.M. Microlaser Technologies AG, Bernried, Germany). Microdissection was performed at 5X magnification. Samples were collected in adhesives caps (P.A.L.M. Microlaser Technologies AG, Bernried, Germany).

cDNA were synthesized from total RNA using RevertAid Premium Reverse Transcriptase (Fermentas) and primed with oligo-dT primers (Fermentas) and random primers (Fermentas). qPCR was performed using a LightCycler^®^ 480 Real-Time PCR System (Roche, Meylan, France). The PCR data were exported and analyzed with a software (Gene Expression Analysis Software Environment) developed at the NeuroCentre Magendie. For the determination of the reference gene, the Genorm method was used. Relative expression analysis was corrected for PCR efficiency and normalized against two reference genes: the peptidylprolyl isomerase A (Ppia) and non-POU-domain-containing, octamer binding protein (Nono). The relative level of expression was calculated using the comparative (2-ΔΔCT) method. Primers sequences are reported below.

**Supplementary Table 1.**
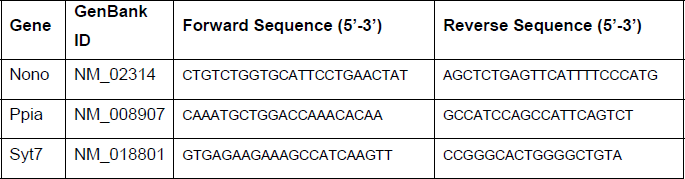
Mouse qPCR primer sequences.

### Cell fractionation in iodixanol gradient

Neuronal cultures were homogenized in buffer A (130 mM KCl, 25 mM NaCl, 25 mM TrisHCl pH 7.4, 1 mM EGTA, protease inhibitors) and centrifuged at 1000g, 10min. The supernatant S1 was then transferred to gradient of iodixanol in buffer A and centrifuged at 122 000 × g, 40 min. Each fractions was then diluted in buffer A and centrifuged at 126 000 × g, 40 min. Pellets were suspended in lysis buffer and analyzed by western-blotting.

### Co-immunoprecipitation and western-blotting

Analysis of the expression of the different polypeptides expressed by lentiviral constructs was tested by transfecting HEK293 cells with the non-liposomal reagent X-tremeGENE (Roche). Cells were scraped and lysed two days after transfection. Analysis of the expression level and the interaction of endogenous neuronal proteins were made from lysates of cortical neurons cultures for 9 DIV in Neurobasal (Gibco) and then up to 14DIV in BrainPhys. The γ-secretase inhibitor (L685,458) was applied from 9 DIV up to lysis. Cells were lysed in a dodecylmaltoside based lysis-buffer (Hepes 50 mM, pH7.4; NaCl 100 mM, Glycerol 10%, DDM 0.5%). Immunoprecipitation was performed using an antibody against Syt7 on 600 to 1000 μg of cell lysates. Immunoglobulins-proteins complexes were pull down by addition of magnetic beads (surebeads) and magnets. Samples were completed with Laemmli buffer. Proteins in samples were separated by western-blotting using 4-20% gradient Tris-tricine gels. Membranes were probed for the proteins of interest and chemio-luminescent bands detected with a CCD camera (ChemiDoc – BioRad).

### Statistical analysis and graphical representations

Statistical analyses were performed with Prism 6.0 (GraphPad Software, La Jolla, CA, USA). Values were first tested for normality (D’Agostino & Pearson omnibus test). Data were presented as scatter plots with mean or median according to the result of the normality test. Normally distributed data were compared using a t-test while Mann-Whitney rank test was used for non-normal data set. Analyses involving two factors (genotype, stimulus number) were presented as points with error bars (± SEM) and non-linear fits. Statistical differences were tested using two-way ANOVAs. Cumulative frequency distributions were presented as fractions and tested using a Mann-Whitney rank test. Statistical differences were considered as significant at p<0.05.

### Drugs

All drugs were obtained from Tocris biosciences or Sigma-Aldrich.

### SUPPLEMENTARY FIGURES LEGENDS

**Supplementary figure 1.**
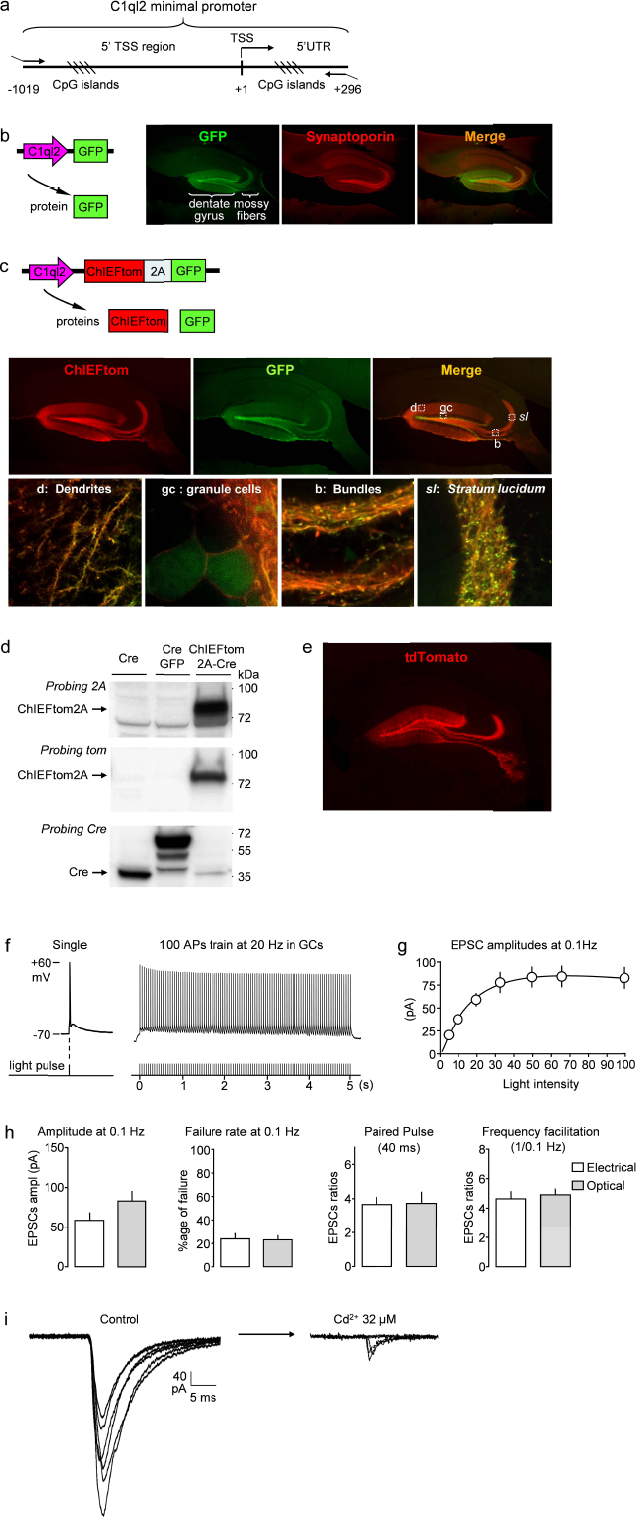
An optogenetic approach to combine cell-specific activation with gene deletion in dentate gyrus granule cells. (**a**) Schematic representation of the mouse C1ql2 minimal promoter cloned from genomic DNA. The 1315 bp sequence includes a region proximal to the transcription start site (5′ TSS) and part of the 5′ untran slated region of the C1ql2 mRNA (5′ UTR). The sequence is rich in putative transcription factor binding sites (CpG islands). (**b**) Expression of GFP controlled by the C1ql2 promoter targets mature Mfs labelled by an antibody against synaptoporin. (**c***)Top panels*, scheme of a bicistronic DNA construct that allows the co-expression of ChIEFtom and GFP under the control of the C1ql2 promoter. *Middle panels*, pictures showing the co-expression of ChIEFtom and GFP in the DG. Small squares indicated in the merge panel are presented enlarged in the lower row of panels. *Lower panels*, images acquired using a confocal microscope with a 20x or 60x objective showing that the two proteins ChIEFtom and GFP are well expressed in all cell compartments (dendrites, GCs soma, Mf bundles, Mf boutons in the stratum lucidum (SL). The fluorescence of mTomato fused to membrane bound ChR2 is prominent in dendrites and fibers whereas the fluorescence of cytosolic GFP is prominent in soma. The two polypeptides are well separated as shown by the expression of the membrane-bound ChIEFtom at the GC cell surface and of the free GFP in the cytosol (panel “gc: granule cells). However, infected neurons do not display any sign of degeneration, toxicity or aggregates of ChR2. (**d**) Western blots of ChIEFtom2A-Cre constructs (right) showing that ChIEFtom is fused to the remaining 2A peptide while Cre recombinase is free. (**e**) Expression of Cre recombinase under the control of the C1ql2 promoter allows the recombination of floxed-tdTomato gene in Ai9 mouse line leading to the expression of free cytosolic tdTomato in GC soma. (**f**) Electrophysiological properties of ChIEF-expressing GCs studied in the whole-cell current clamp mode. *Left*, a single action potential (AP) evoked by a light pulse at 470 nm of 1 ms. *Right*, a long burst of 100 stimuli at 20 Hz for 5 s triggers a long train of APs without failure. (**g**) Plot of EPSC amplitudes in function of light intensity recorded from 20 CA3 pyramidal cells. 1 ms pulses of 470 nm light were shinned on top of GC dendrites using a 63x objective. Light intensity was increased up to 2 mW/cm2. (**h**) Comparison of averaged values of Mf-EPSC amplitude, failure rate, paired-pulse and frequency facilitation recorded in CA3 pyramidal cells when stimulating Mfs with an extracellular electrode placed in the hilus or by stimulating GC dendrites with light. (**i**) Representative Mf-EPSCs recorded in CA3 pyramidal cells in control conditions or in presence of 32 μM Cadmium^2+^ (Cd^2+^). The presence of Cd^2+^ dramatically increases the failure rate and decreases the amplitude of the >EPSCs.

**Supplementary figure 2.**
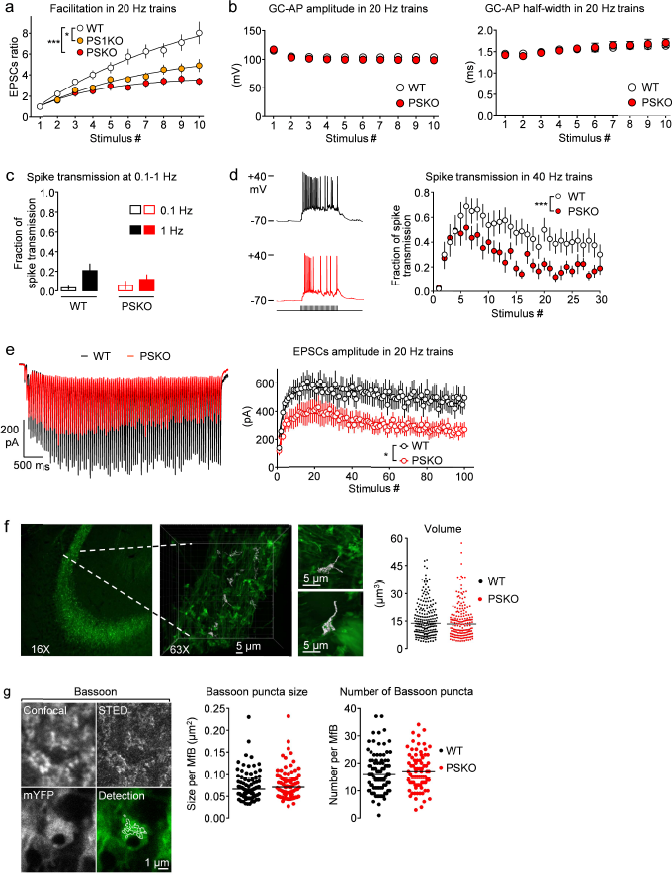
Electrophysiological properties of GCs, Mf-CA3 neurotransmission and Mf boutons morphology. (**a**) Analysis of Mf-EPSCs recorded in CA3 pyramidal cells while stimulating with ten repetitive light pulses at 20 Hz in WT, PS1KO and PS1/PS2KO conditions. Facilitation is represented by the ratio of EPSC amplitudes relative to the amplitude of the EPSC evoked by the first stimulus. Note that facilitation is impaired in the absence of PS1 alone. Two-way ANOVA, *p=0.0304.* (**b**) *Left*, Plot of AP amplitudes in GCs soma recorded in the current-clamp mode evoked by 10 light pulses of 1 ms at 20 Hz on GCs in WT and PSKO conditions. *Right*, Plot of APs half-width in conditions as in *“Left”.* (**c**) *Left*, histograms of the mean fraction of spike transmitted in CA3 pyramidal cells recorded in the current-clamp mode at 0.1 or 1 Hz in WT and PSKO conditions. Note that in average the fraction of spike transmitted is low in all conditions even if a few neurons do fire quite reliably at 1 Hz. (**d**) *Left*, Representative traces of APs in CA3 pyramidal cells in response to the stimulation of GC dendrites at 40 Hz in WT (black) and PSKO (red) conditions. *Right*, plot of the fractions of spike transmission (for 16 recorded neurons per genotype; 5 sweeps per neuron) in WT or PSKO genotype. Note that at 40 Hz the highest rate of transmission is reached around 5 stimuli and that the plot in PSKO is lower than in WT condition (Two-way ANOVA, *p<0.0001).* (**e**) *Left*, representative Mf EPSCs obtained by illuminating the dendrites of GCs with 100 stimuli at 20 Hz. After the peak reached within ten stimuli, the amplitudes of Mf-EPSCs decrease while the charge of the train increases. Right, analysis of several recordings obtained as in *“Left”.* The raw amplitude of Mf-EPSCs is plotted against the stimulus number. Note that Mf-EPSC amplitudes in PSKO condition are lower than in WT. Two-way ANOVA, n=20/genotype, *p<0.05.* (**f**) *Left*, confocal images of the *stratum lucidum* region of the hippocampus in brain slices where the Mfs projects onto CA3 pyramidal cells. *Middle*, zoom in using a 63x objective allows to observe Mf boutons. 3D reconstruction of some Mf boutons after acquisition of Z-stack are depicted in grey. A zoom on two large Mf boutons allows to appreciate their large volume and complex structure. *Right*, scatter plots with medians of the volume of Mf boutons in WT and PSKO conditions. (**g**) Evaluation of Bassoon puncta number and surface using STED microscopy. *Left*, image of a Mf bouton labelled with membrane-bound YFP (mYFP, bottom left panel) and stained against Bassoon (upper left panel in confocal; upper right panel in STED microscopy). Detected Bassoon positive puncta are delimited with white lines. *Middle:* scatter plots of average size of Bassoon puncta per Mf bouton. *Right*, scatter plots of average number of Bassoon puncta per MfB. 76 Mf boutons have been analyzed per genotype. The absence of PS does not impact on the number of putative releasing sites.

**Supplementary figure 3.**
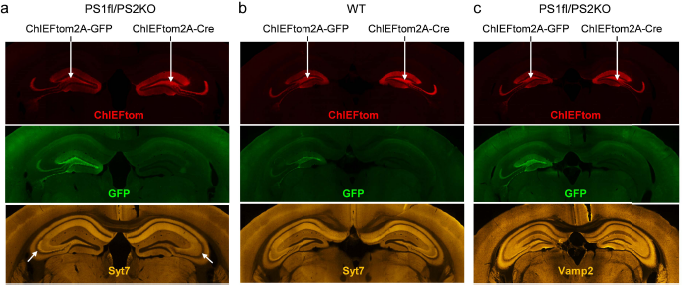
Effect of targeted deletion of PS in GCs on Syt7 and Vamp2 expression. (**a-c**) Pictures of coronal sections from mice injected in the left hemisphere with a LV expressing ChIEFtom and GFP and on the right hemisphere with a LV expressing ChIEFtom and Cre. ***(a)*,** in PSfl1/PS2KO mice, a decrease of Syt7 staining is observed only in the SL where PS1 has been removed (compare the two oblique arrows). In WT condition stained for Syt7 ***(b)*** or in PSfl1/PS2KO mice stained for Vamp2 ***(c)*,** no difference is observed between the two hemispheres. Note that the absence of PS2 alone does not noticeably affect the staining of Syt7 compared to WT (compare the left hemispheres of WT and PS1fl/PS2KO conditions).

**Supplementary figure 4.**
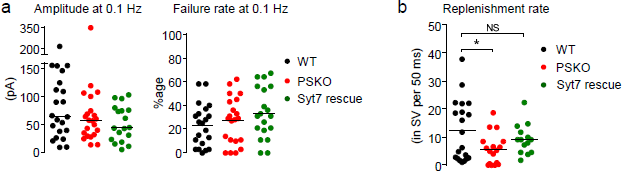
Basal transmission at PSKO Mf-CA3 synapses rescued for Syt7. (**a**) Basal transmission at Mf-CA3 synapses in WT, PSKO and Syt7-PSKO conditions. *Left*, scatter plots with medians of EPSCs amplitude recorded in CA3 pyramidal cells in the whole cell voltage-clamp mode while stimulating GCs with light at 0.1 Hz. *Right*, scatter plots with mean failure rates. (**b**) Scatter plots with means of replenishment rates at 40 Hz calculated neuron per neuron are represented by genotype. The replenishment rate, impaired in the absence of PSKO, is rescued by re-expression of Syt7.

**Supplementary figure 5.**
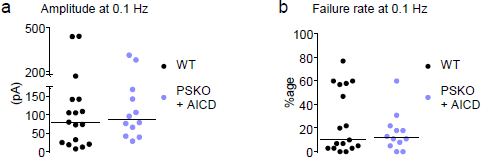
Basal transmission at PSKO Mf-CA3 synapses with rescue of AICD. (**a**) Basal transmission at Mf-CA3 synapses in WT and PSKO with re-expression of AICD. Scatter plots with medians of EPSCs amplitude recorded in CA3 pyramidal cells in the whole cell voltage-clamp mode while stimulating GCs with light at 0.1 Hz. (**b**) Scatter plots with mean failure rates in conditions as in “a”.

**Supplementary figure 6.**
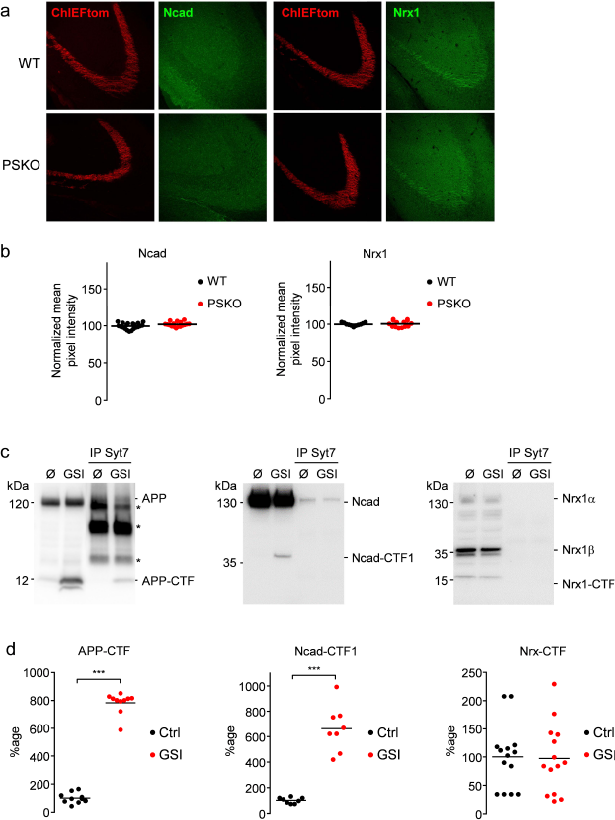
Expression of γ-secretase substrate candidates in Mfs and accumulation of C-terminal stubs upon γ-secretase inhibition. (**a**) Confocal images of the Mfs tracts in WT and PSKO conditions observable in the red channel (ChIEFtom) and green channel (staining for Ncadherin Cterminal (Ncad) or Neurexin Cterminal (Nrx1)). Mfs are faintly labelled by the Ncad antibody and more prominently by the Nrx1 antibody. The absence of PS does not alter labelling with these two antibodies. (**b**) Scatter plots of the mean pixel intensity in the *SL* normalized to the mean pixel intensity in the *stratum radiatum* for Ncad and Nrx1 labelling. The absence of PS does not increase the intensity of the labelling indicating that Cterminal stubs of Ncad or Nrx1 do not accumulate in Mfs. (**c**) Western-blot of protein extracts from neuronal cultures treated or not with a γ-secretase inhibitor (GSI). GSI induces the accumulation of APP-CTFs and Ncad Cterminal stub (Ncad-CTF1) but not of the band expected to be Neurexin1-CTF (16 kDa). IP of Syt7 pull down APP-FL, βCTF and Ncad but not Ncad-CTF1 or Neurexin1 of any form. (**d**) Quantitative analysis of the CTF bands detected in western blot of starting materials from neuronal cultures treated or not with GSI as in “c”. An increase of 800% in the expression of APP-CTF was detected. An increase of 650% in the expression of Ncad-CTF was detected. The expression level of Neurexin-CTF was unchanged by treatment with GSI.

